# Synergistic interference with SARS-CoV-2 replication by Molnupiravir-derived N4 hydroxycytidine and inhibitors of CTP synthetase

**DOI:** 10.1101/2025.03.06.641790

**Authors:** Kim M. Stegmann, Antje Dickmanns, Hannah L. Fuchs, David Scheibner, Björn-Patrick Mohl, Florian R. Moeselaken, Wencke Reineking, Theresa Störk, Asisa Volz, Philip A. Beer, Andrew E. Parker, Veronika Pilchova, Christian Meyer zu Natrup, Maren von Köckritz-Blickwede, Wolfgang Baumgärtner, Anne Balkema-Buschmann, Matthias Dobbelstein

## Abstract

N4-hydroxycytidine (NHC), the active metabolite of Molnupiravir, is incorporated into nascent RNA of SARS-CoV-2 and interferes with subsequent virus replication. We have previously described synergy between NHC and inhibitors of dehydroorotate dehydrogenase (DHODH), an enzyme required for pyrimidine synthesis. Upon DHODH inhibition, the lack of endogenous pyrimidines conceivably enhances NHC incorporation. However, the question remains whether preventing the synthesis of just one pyrimidine base, cytidine, might as well augment the antiviral efficacy of NHC. We tested this by inhibiting CTP synthetases (CTPSs), the cellular enzymes that directly catalyze the synthesis of a cytidine nucleotide. We observed that inhibitors of CTP synthetase (CTPSis), namely cyclopentenyl cytosine (CPEC) as well as STP938 and STP720, display a strong synergy with NHC for diminishing SARS-CoV-2 replication in cell culture, as shown earlier for DHODH inhibitors. NHC and CTPSis in combination prevented the cytopathic effect of SARS-CoV-2 and strongly reduced the release of viral RNA and infectious particles, as well as the synthesis of viral proteins. This combination was also active against an Omicron variant of SARS-CoV-2. Addition of cytidine, but not uridine, rescued virus growth under these conditions. Of note, treating SARS-CoV-2-infected hamsters with the CTPS1 inhibitor STP938 strongly diminished COVID pathology. We propose that CTPS inhibition has the potential to increase the efficacy of antiviral cytidine analogues and to treat coronavirus infections.

**HIGHLIGHTS:**

- The efficacy of NHC against SARS-CoV-2 replication in cell culture models is intensified by several orders of magnitude through targeting cellular CTP-Synthetase.
- The drug combination still displays its effect against SARS-CoV-2 replication in the presence of uridine, suggesting that serum uridine cannot counteract its efficacy.
- CTPS inhibition diminishes COVID-19-like pathology in an established animal model.

## INTRODUCTION

The COVID pandemic has caused millions of deaths, and new waves of infections caused by SARS-CoV-2 or related viruses may rise in the future. This illustrates the need for broadly active and efficient antiviral therapies (Chan et al., 2024; Steiner et al., 2024). One treatment option against COVID-19 consists in an antiviral nucleoside analogue termed Molnupiravir. The drug was authorized or approved for clinical use in several countries (Maas et al., 2024), although this was withdrawn or replaced by emergency-only approval more recently. Molnupiravir is orally available and is metabolized to N4-hydroxycytidine (NHC) in the liver. Upon cellular uptake, NHC is triphosphorylated and, when the cell is infected by SARS-CoV-2, incorporated into nascent viral RNA (Toots et al., 2019). This causes ambiguous base pairing in subsequent rounds of replication, since NHC not only forms base pairs with the canonical guanine, but also with adenine (Kabinger et al., 2021). As a result, the virus undergoes extensive mutagenesis, diminishing its viability and explaining the therapeutic efficacy of Molnupiravir. Despite being mutagenic to the viral RNA (Kumar et al., 2024; Zibat et al., 2023), the drug hardly gives rise to virus mutants resistant against it, and it is broadly active against variants of SARS-CoV-2 (Strizki et al., 2024).

In previous work, we and others have reported a strategy for further increasing the therapeutic impact of this drug, by combining it with inhibitors of dihydroorotate dehydrogenase (DHODH) (Schultz et al., 2022; Stegmann et al., 2022). In addition to NHC, DHODH inhibition also cooperates with the nucleoside analogue 4’-fluorouridine against a panel of RNA viruses (Schrell et al., 2025). DHODH inhibitors (DHODHis) reduce the amount of available pyridimidine nucleotides in the cell, enabling their clinical use against cancer (Mollick and Laín, 2020), autoimmunity (Leban and Vitt, 2011), and infections (Boschi et al., 2019). DHODH inhibition thus increases the likelihood of incorporating NHC during virus replication, conceivably by decreasing the intracellular levels of pyrimidine nucleotides that would otherwise compete with NHC for incorporation. The combination of DHODHis with NHC reduces virus replication by several orders of magnitude in comparison to the single drugs (Stegmann et al., 2022). However, when we tried the same combination to treat SARS-CoV-2-infected hamsters, this only moderately decreased virus load and disease progression (Stegmann et al., 2022). As a possible explanation, the sera of most mammals contain 2 to 8 µM uridine (Karle et al., 1980) in a tightly regulated fashion (Deng et al., 2017). We showed that similar concentrations were sufficient to rescue virus replication in vitro, despite the presence of both drugs (Stegmann et al., 2022). We therefore propose that the uptake of extracellular uridine and its subsequent phosphorylation may compensate for the lack of endogenous pyrimidine synthesis when applying DHODHis. The available uridine nucleotides would then serve as substrates for the synthesis of cytidine nucleotides, and this would compete with NHC nucleotides for incorporation into virus RNA, thereby diminishing the efficacy of the drug combination in vivo.

To improve the therapeutic impact of NHC/Molnupiravir, we investigated an alternative strategy of interfering with pyrimidine synthesis. Rather than reducing the available uridine through DHODHis, we now inhibited cytidine triphosphate (CTP) synthetase (CTPS; the name ‘synthetase’ traditionally describes an enzyme that hydrolyses ATP or another nucleoside triphosphate, as is the case for CTPS; however, the broader term ‘synthase’ is also widely accepted to describe CTPS). CTPS is responsible for the synthesis of CTP from uridine triphosphate (UTP), transferring a -NH_2_ moiety from glutamine to UTP. Thus, the inhibition of CTPS cannot be overcome by the cellular uptake of uridine. In humans, the cytidine serum levels are considerably lower than the uridine levels (Cansev, 2006). Thus, nucleotide salvage by extracellular cytidine is less likely to overcome the impact of CTPS inhibition, as opposed to the extracellular uridine that antagonizes the efficacy of DHODHis. With these considerations, we tested whether CTPS inhibitors (CTPSis) are capable of improving the antiviral effect of NHC/Molnupiravir.

In earlier studies, CPTS inhibition was achieved by cyclopentenyl cytosine (CPEC) (Schimmel et al., 2007). At that time, this drug was mostly employed as an antineoplastic agent with promising initial results. However, its intravenous infusion into patients at high doses resulted in fatal cases of unexplained hypotension (Politi et al., 1995). This led to discontinuation of investigating CPEC in clinical studies.

In the meantime, however, improved CTPSis were designed for treating autoimmune (Soudais et al., 2024) or malignant (Asnagli et al., 2023; Pfeiffer et al., 2024) diseases. Whereas CPEC requires intracellular phosphorylation for efficacy, the novel CTPSis are active in their native forms (Lynch et al., 2021).

Here we show that CTPSis strongly synergize with NHC/Molnupiravir in interfering with the replication of SARS-CoV-2 in vitro, with a degree of synergy that resembles DHODHis. The addition of cytidine but not uridine rescues virus replication in the presence of CTPSis. Moreover, CTPSis diminished early virus replication and lung pathology in a hamster model of COVID-19.

## RESULTS

### CTPSis synergize with NHC to reduce the SARS-CoV-2-induced cytopathic effect

Similar to our previous work, we combined NHC with inhibitors of pyrimidine synthesis. The idea behind this approach is that reduced pyrimidine nucleotides will increase the chance of incorporating NHC nucleotides into virus RNA during replication. However, instead of inhibiting DHODH (Stegmann et al., 2022), we now specifically reduced the synthesis of the cytidine nucleotide CTP using CPEC or the CTPSis STP938 and STP720. STP938 has a higher potency against the isoenzyme CTPS1 than CTPS2 (Asnagli et al., 2023), and it is currently being used in clinical studies for treating B and T cell lymphomas (NCT05463263) or solid tumors (NCT06297525). STP720 inhibits both CTPS isoenzymes to similar degrees. We expected that CTPS inhibition should reduce the levels of available cytidine nucleotides and thus increase NHC incorporation (Fig. 1A).

**Figure 1:**
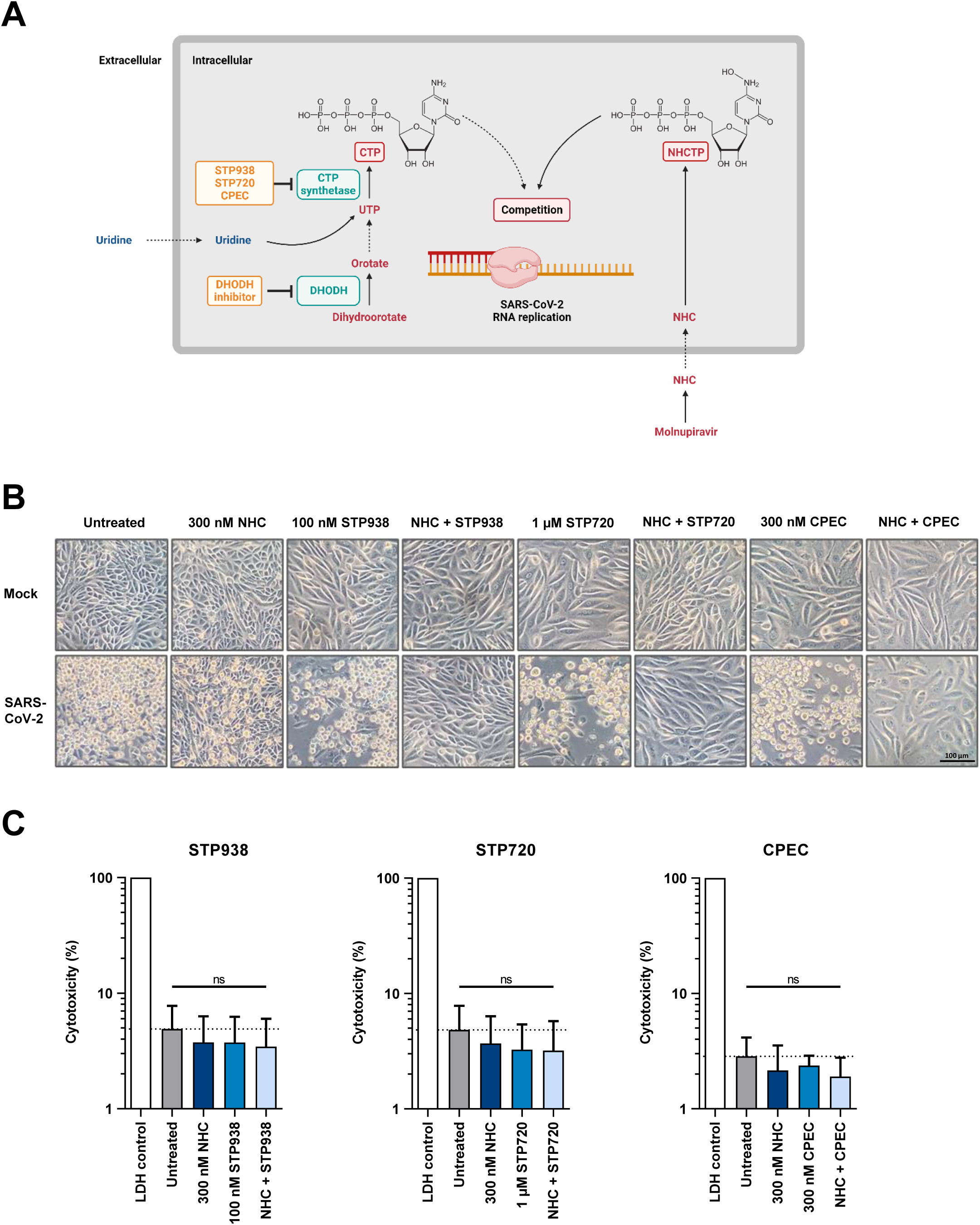
Diminished SARS-CoV-2-associated Cytopathic Effect (CPE) in the presence of CTPS inhibitors and NHC. A. Metabolic pathways explaining the therapeutic rational. Pyrimidine synthesis first yields uridine triphosphate (UTP), and the transfer of an amino group by cytidine triphosphate (CTP) synthetase (CTPS) results in CTP. CTP directly competes with N4-hydroxycytidine (NHC) triphosphate (NHCTP), derived from the Molnupiravir-metabolite NHC, for incorporation into the genomic RNA of SARS-CoV-2. Thus, inhibition of CTPS facilitates the incorporation of NHC nucleotides and therefore enhances the efficacy of Molnupiravir/NHC. B. CTPS inhibition and NHC jointly diminish the SARS-CoV-2-driven CPE. Vero E6 cells were treated with 300 nM cyclopentenyl cytosine (CPEC; targeting both isoenzymes of CTPS, CTPS1 and CTPS2) or 100 nM STP938 (preferentially targeting CTPS1), or 1 µM STP720 (targeting CTPS1 and CTPS2), alone or in combination with 300 nM NHC as indicated. After 24 hrs, the cells were infected with SARS-CoV-2 and further incubated with the drugs for another 48 hrs. Cell morphology was documented by phase contrast microscopy to reveal that the CPE was diminished with the drug combinations. Bar, 100 µm. C. Lack of overt cytotoxicity by the drugs, based on LDH release. Lactate dehydrogenase (LDH) in the supernatant of drug-treated cells was quantified as a proxy for cytotoxicity. The drugs did not markedly affect the amount of released LDH, indicating lack of cytotoxic cell lysis (mean with SD, n=3). For cytotoxicity data with higher doses of CTPSis, see Suppl. Fig. 1.

We treated Vero E6 cells with CTPSis and/or NHC, followed by infection with SARS-CoV-2. The drugs alone did not grossly affect cell integrity, as determined by morphology (Fig. 1B) and release of lactate dehydrogenase (LDH; Fig. 1C, Suppl. Fig. 1), although we did observe reduced cell proliferation when cells were exposed to CTPSis for several days – most likely as a result of diminished nucleotide levels. A cytopathic effect (CPE) was extensively induced by SARS-CoV-2 in non-treated cells. Both CTPSis or NHC alone only slightly reduced the CPE. Importantly, however, the combination of CTPSis and NHC completely eliminated any visible CPE (Fig. 1B), strongly suggesting that they were together abolishing the propagation of SARS-CoV-2 in treated cells.

### CTPSis cooperate with NHC to reduce SARS-CoV-2 replication

To assess SARS-CoV-2 replication directly, we prepared RNA from the supernatants of infected cells and quantified the amounts of virus-derived RNA by qRT-PCR. Strikingly, the combination of CTPSis and NHC reduced the amount of virus RNA derived from Vero E6 cells by several orders of magnitude, with drug concentrations that were at best moderately affecting virus replication when used alone (Fig. 2A, Suppl. Fig. 2A-C, Suppl. Table 1). The high synergy scores that we derived from the RNA quantities (Fig. 2B) also reflected this observation. Similar synergy was also observed when treating and infecting Calu-3 cells in an analogous manner (Fig. 2C,D, Suppl. Fig. 2D). Taken together, these observations indicate synergistic antiviral activity of CTPSis and NHC in cell culture models.

**Figure 2:**
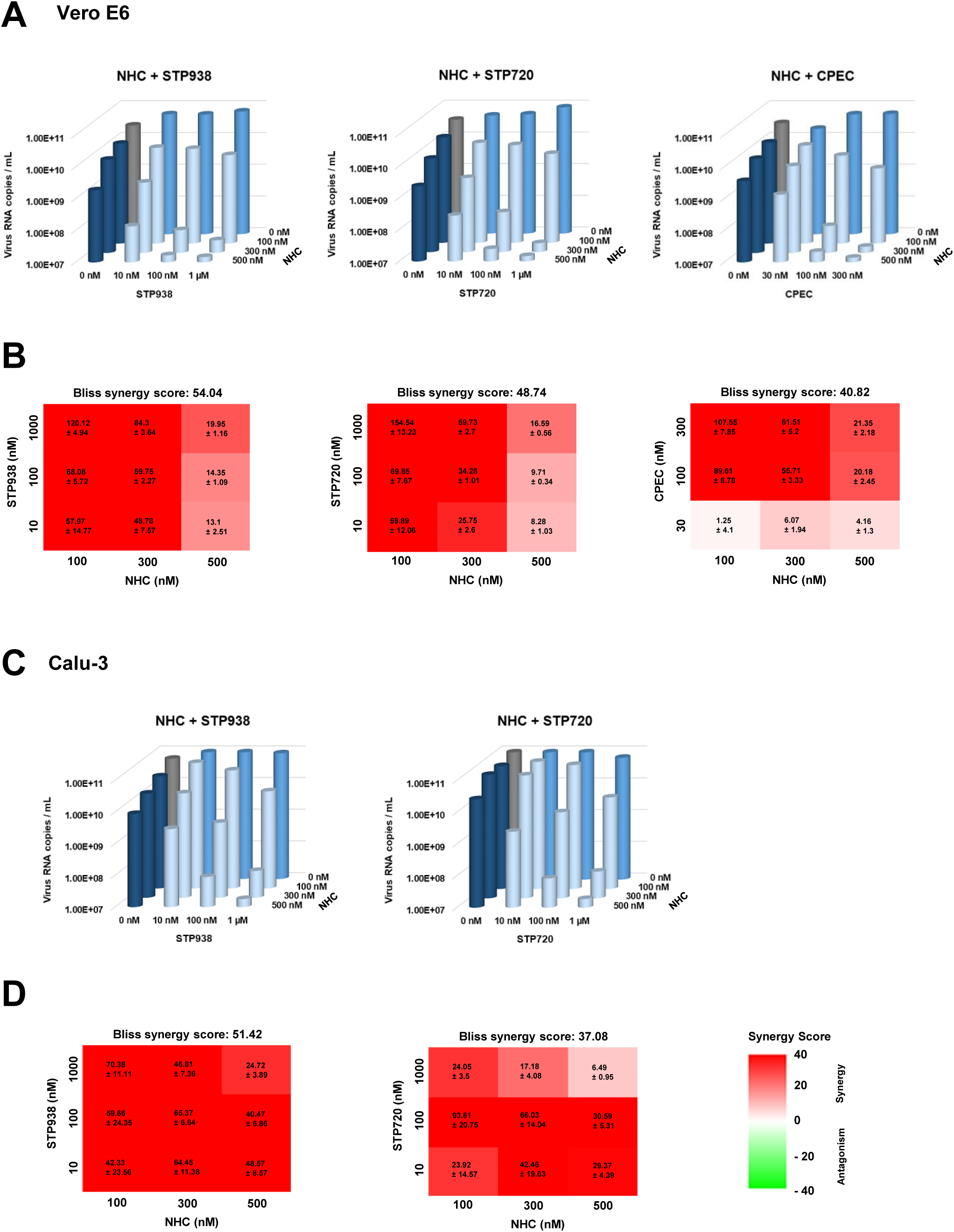
Synergistic suppression of virus production upon cell treatment with CTPSis and NHC. A. Viral RNA in the supernatant of treated/infected cells. Vero E6 cells were treated with CTPSis and/or NHC and infected with SARS-CoV-2 as in Fig. 1B, followed by quantification of virus RNA in the supernatant by RT-PCR, indicating synergistic antiviral effects of both drugs (mean, n=3). For single values of each replicate, see Suppl. Fig. 2A-C. For p-values, see Suppl. Table 1. B. Synergy scores revealed strongly synergistic antiviral activities of CTPSis and NHC. The synergy scores were calculated using the Bliss independence model. Data are presented as mean ± SEM. A Bliss score > 10 is generally considered to reveal strong drug synergism (n=3). C. Analogous experiments as in A were carried out using Calu-3 cells (n=3). D. Analogous evaluations as in B were carried out using Calu-3 cells (n=3).

### The drugs synergize to inhibit virus protein synthesis and virus propagation

To further corroborate the observed drug synergy, we visualized viral proteins in SARS-CoV-2 infected and treated Vero cells by immunofluorescence (Fig. 3A-C) and immunoblot (Fig. 3D-F) analyses. Neither CTPSis nor NHC were capable of reducing the amount of the viral Nucleocapsid (N) or Spike (S) proteins in the applied concentrations. However, when combined, both proteins essentially disappeared, again confirming the strong antiviral synergy of the drugs.

**Figure 3:**
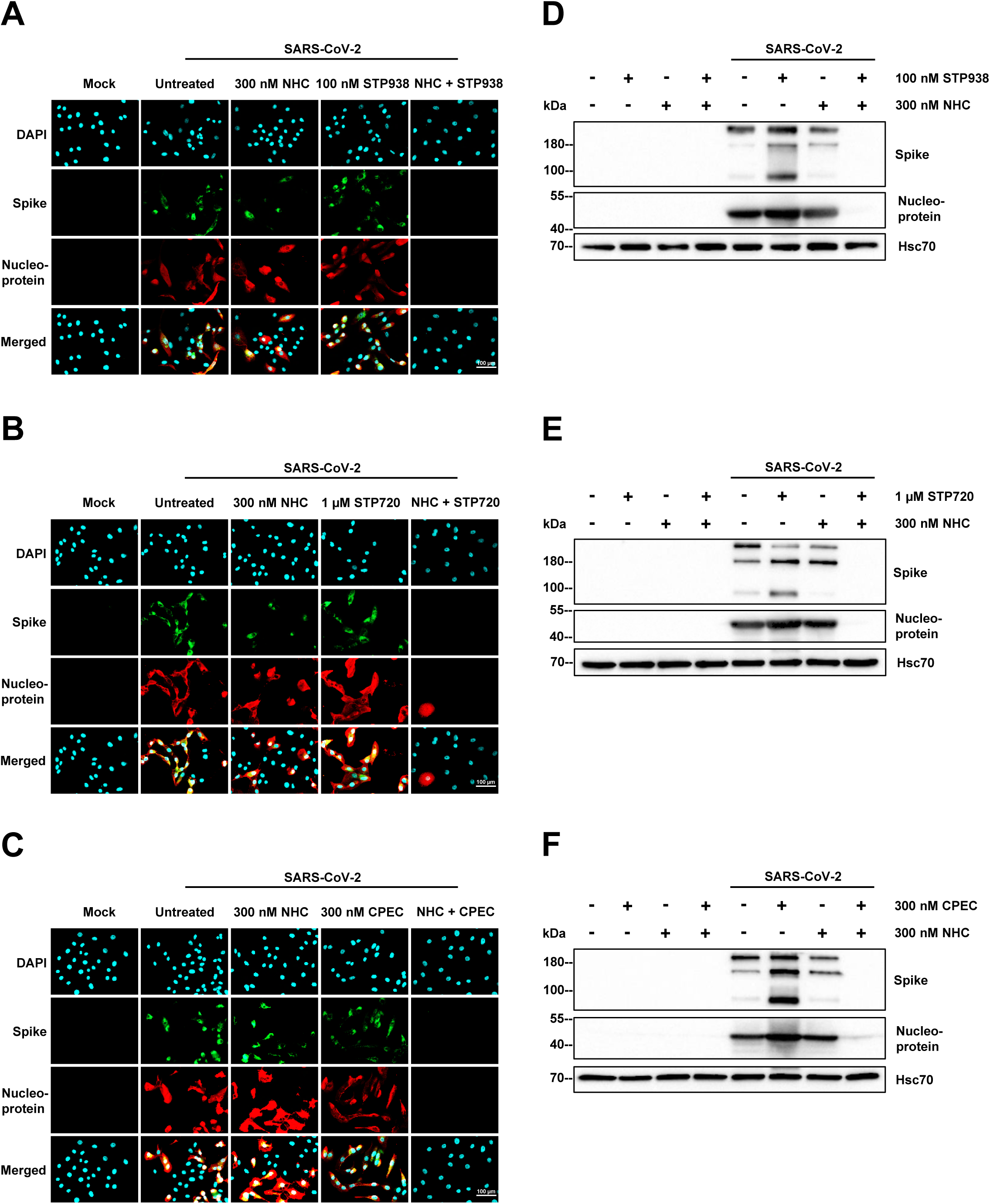
Strongly reduced synthesis of virus proteins upon combined drug treatment. (A-C) Immunofluorescence analysis. Vero E6 cells were treated with NHC and/or STP938 (A), STP720 (B), or CPEC (C), and infected as in Figs. 1 and 2; twenty-four hours after infection, the SARS-CoV-2 nucleoprotein (N) and spike protein (S) were detected by immunofluorescence microscopy. (D-F) Upon treatment and infection as in A, the cells were harvested and subjected to SDS-PAGE and immunoblot analysis to detect N and S, indicating strong reduction by the drug combination but not by the single drugs at the same dose.

To analyze the production of infectious particles, we applied CTPSis and NHC at concentrations that only moderately suppressed virus replication when used as single treatments, infected cells with SARS-CoV-2 and determined the median tissue culture infectious dose (TCID_50_/mL) of the supernatant (Fig. 4A). Remarkably, the combination of CTPSis and NHC was far more efficient in reducing the TCID_50_/mL as compared to untreated samples or treatments with single drugs. Moreover, the combination treatment reduced the virus RNA progeny even when applied 4 hrs after infecting the cells with SARS-CoV-2 (Fig. 4B). Furthermore, we evaluated the replication of the SARS-CoV-2 Omicron variant B.1.1.529 in the presence of CTPSis and NHC (Fig. 4C). Our findings confirm that the drug combination effectively combats this highly mutated variant as well.

**Figure 4:**
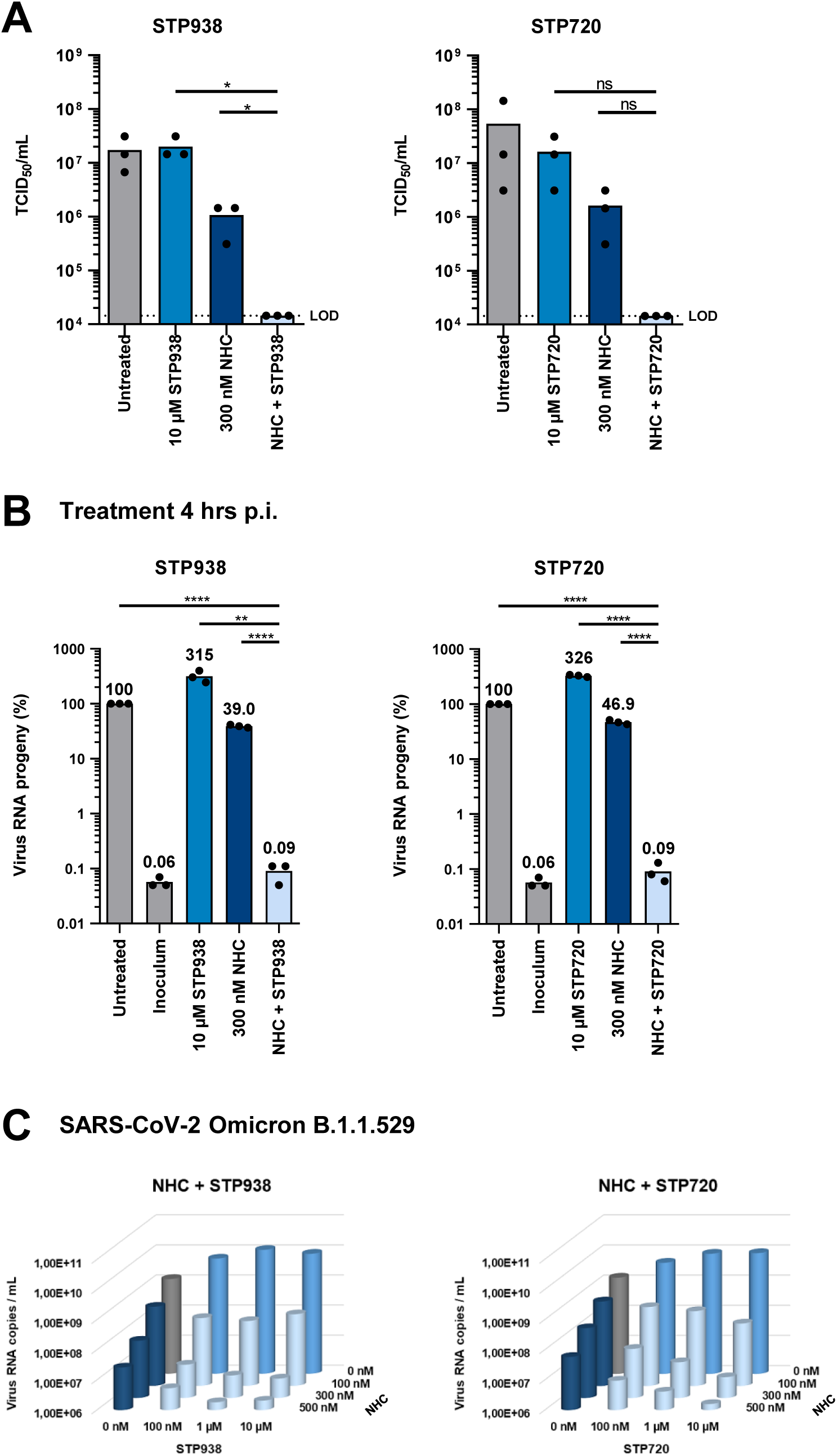
Drug synergy addressed by additional approaches. A. TCID50 revealing drug synergism. After treatment and infection as in Figs. 1 and 2, the infectious virus particles within the supernatant were quantified as Tissue Culture Infectious Dose 50 (TCID50/mL), largely corresponding to the quantification of virus RNA (mean with single data points, n=3). Statistical significance is indicated by asterisks as follows: **** for p ≤ 0.0001; *** for p ≤ 0.005; ** for p ≤ 0.01, * for p ≤ 0.05 and ns for not significant (p > 0.05). B. Virus RNA progeny was diminished by CTPSis and NHC even when the drugs were added only 4 hrs post infection (p.i.). Vero E6 cells were infected with SARS-CoV-2 as described in Figs. 1 and 2 and treated with CTPSis and/or NHC 4 hrs later. RNA was extracted from the cell culture supernatant 48 hrs later, to quantify SARS-CoV-2 RNA progeny by qRT-PCR. The RNA amounts obtained upon infection without drug treatment served as the baseline at 100%, and the other RNA quantities were normalized accordingly (mean with single data points, n = 3). Statistical significance is indicated by asterisks as in A. C. The combination of CTPSis with NHC reduces the replication of the SARS-CoV-2 Omicron variant B.1.1.529. Vero E6 cells were treated as described in Fig. 1B and infected with the SARS-CoV-2 Omicron variant. After 48 hrs of virus infection, the RNA was isolated from the cell supernatants, followed by qRT-PCR to determine the amount SARS-CoV-2 RNA copies per mL (mean, n=3).

### Synergistic action of CTPSis and NHC is rescued by cytidine but hardly by uridine

We were investigating the combination of CTPSis and NHC based on the assumption that CTPSis reduce the amount of intracellular cytidine nucleotides and thus increase the incorporation of NHC nucleotides into virus RNA (Fig. 1A). To scrutinize this concept, we added nucleosides while treating the cells with CTPSis and/or NHC, followed by virus infection. As expected, the addition of cytidine rescued the replication of SARS-CoV-2 despite the presence of CTPSis and NHC, whereas the addition of uridine, even at 50 µM concentration, largely failed to reconstitute virus replication; this was observed using two different CTPSis, by qRT-PCR to quantify virus RNA in the cell supernatant, as well as by immunoblot analysis of virus proteins (Fig. 5A-F). This concentration exceeds that of uridine in the serum of most mammalians by 5 to 15 fold (Yamamoto et al., 2011). These experiments argue that extracellular uridine, at concentrations typically found in serum, should not be able to interfere with the synergistic antiviral effects of CTPSis and NHC.

**Figure 5:**
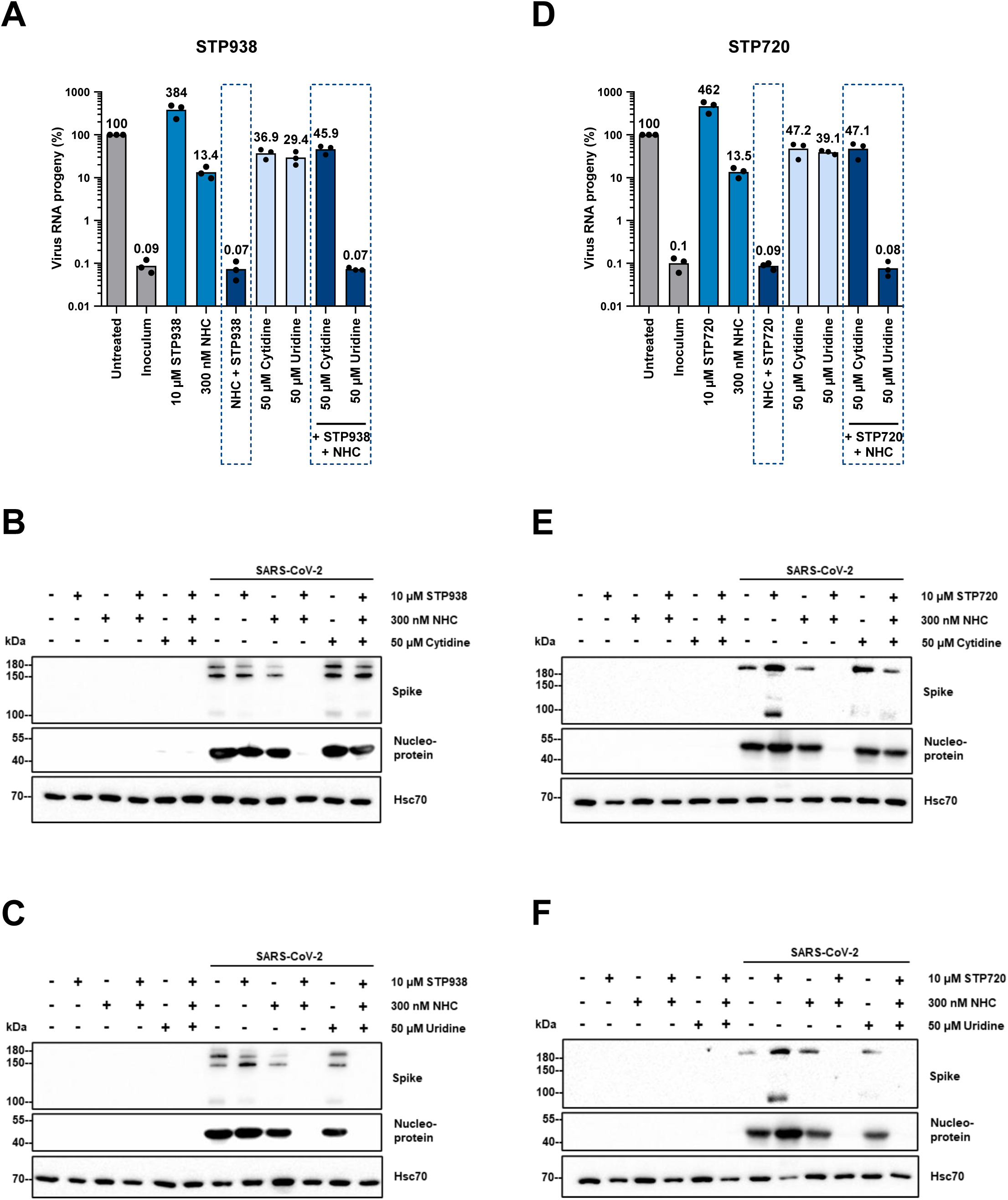
Efficient rescue of virus replication by cytidine but not uridine. (A) Vero E6 cells were treated with STP938 and/or Molnupiravir as described in Fig. 1B, followed by infection and subsequent preparation of virus RNA as in Fig. 2A. RT-PCR analysis revealed synergistic suppression of virus RNA release. In addition to the antiviral treatment, however, we added 50 µM uridine or cytidine. The addition of cytidine led to a rescue of virus RNA levels, whereas the addition of uridine did not (mean with single data points, n=3). (B+C) Vero E6 cells were treated and infected as in (A). In addition to the combination treatment, cytidine (B) or uridine (C) were added. At 48 hrs p.i., cells were harvested and the viral S and N proteins were visualized by immunoblot analysis. Again, cytidine but not uridine rescued the appearance of virus proteins. (D) Analogous experiments as in A were carried out using STP720. (E+F) Analogous experiments as in B+C were carried out using STP720.

### A CTPS1 inhibitor ameliorates COVID-19 pathology in a hamster model

Finally, we sought to determine the efficacy of CTPSis in combination with Molnupiravir, the prodrug of NHC, in an animal model of COVID-19. To limit the number of animals in our proof of concept animal study, we applied one dose using one application route in the Syrian hamster model. We chose the CTPS1 inhibitor STP938 due to its strong in vitro-performance even at low concentration and combined it with Molnupiravir. 24 hrs after the initial treatment, we infected the animals with SARS-CoV-2 and observed the accumulation and shedding of virus as well as tissue pathology (Fig. 6A). Strikingly, STP938 alone reduced the level of shed viral RNA in oral swabs and nasal lavage, in particular at the time points 3 and 4 days post infection (Fig. 6B). These effects were less pronounced when analyzing the amount of infectious particles (Suppl. Fig. 3) by postmortem analysis of the respiratory tract tissues on day 6. Interestingly, Molnupiravir did not further reduce the level of detected viral RNA, but rather restored its production to a moderate extent. While the underlying mechanisms are not fully clear, we speculate that some of the administered Molnupiravir might be metabolized to cytidine in the hamster, which could then compensate for the effect of highly efficient CTPS inhibition by STP938. When investigating the pathologic alterations in bronchial and lung tissues, the COVID-like impact of the virus was strongly attenuated by the administered dose of STP938, and less so upon addition of Molnupiravir (Fig. 6C, Suppl. Figs. 4 and 5). Lung lesions were significantly reduced in the ST938 group compared to the untreated SARS-CoV-2 infected hamsters (p = 0.037). Likewise, the interferon response, as determined by qRT-PCR-detection of *MX1*, was lowest in the cohort treated with STP938 alone (Fig. 6D). In all cases, the combination of Molnupiravir and STP938 was not more effective than the single STP938. Immunohistochemistry (IHC) revealed that the Mx1 protein was present in the cytoplasm of bronchial epithelial cells, vascular endothelial cells, pneumocytes and alveolar macrophages of SARS-CoV-2-infected but untreated animals. In the STP938 only group, immunoreactivity for Mx1 protein was strongly reduced compared to the untreated animals and only slightly higher than in the uninfected control group. In contrast, treatment with Molnupiravir only and especially with the combination resulted in Mx1 IHC scores comparable to the untreated, SARS-CoV-2-infected hamsters (Fig. 6E, Suppl. Fig. 6). The Mx1 immunoreactivity in the STP938 group was significantly reduced compared to untreated SARS-CoV-2-infected hamsters (p=0.0104) and the combined treatment group (p=0.0087).

**Figure 6:**
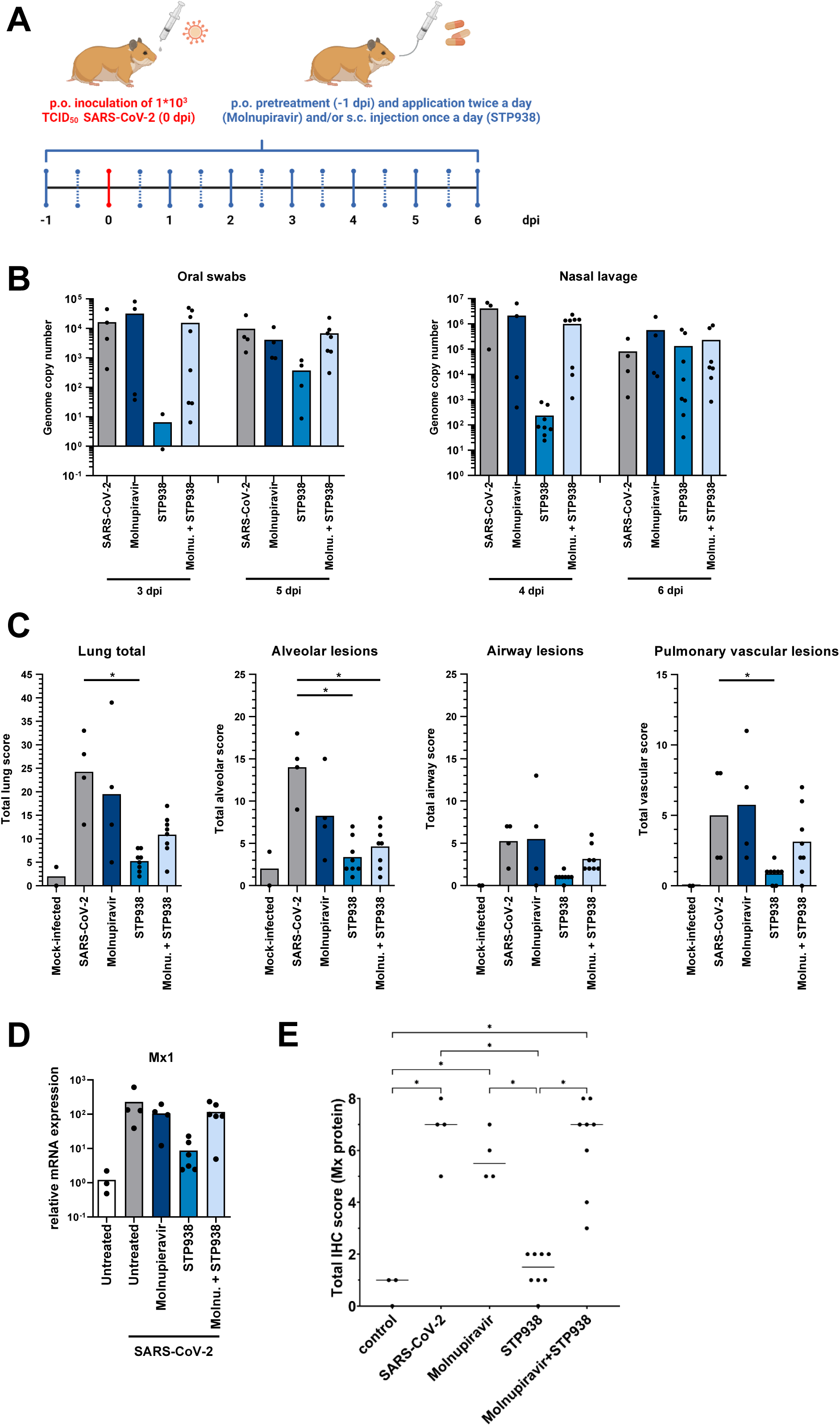
Therapeutic efficacy of STP938 and Molnupiravir in the COVID-19 hamster model. A. Treatment and infection scheme. Hamsters were treated with STP938 and/or Molnupiravir 24 hrs pre infection, followed by orotracheal infection with SARS-CoV-2 and continued treatment. Oral swabs and nasal lavage samples were alternately collected every other day. At day 6 post infection, animals were sacrificed to analyze virus load and tissue morphology in various organs. B. Virus load in oral swabs and nasal lavages was found reduced by STP938 treatment at early time points (day 3-5), as revealed by RT-PCR analysis of virus RNA within the samples. C. Histopathological analysis of lungs, alveolar lesions and pulmonary vascular lesions (H&E stains). STP938 decreased the total lung score (total sum of alveolar score, airway score and pulmonary vascular lesions) compared to the non-treated but infected group. Statistical evaluation was performed by the Kruskal-Wallis test with Benjamini Hochberg correction for multiple comparison (FDR 0.05). While alveolar lesions improved in all treatment groups, vascular lesions were more pronounced in the Molnupiravir treated group compared to the non-treated SARS-CoV-2 group. Molnupiravir also enhanced vascular lesions in the synergistically treated group compared to the STP938 treated group, but overall less pronounced than in the untreated group. For representative images of hamster tissues, see Suppl. Fig. 4. D. Analysis of Mx1 mRNA as a read-out for the interferon response. RNA samples derived from hamster lungs at day 6 post infection were analyzed by qRT-PCR to assess the interferon response in animals upon infection and treatment with Molnupiravir and/or STP938. The mRNA encoding the interferon-induced GTP-binding protein Mx1 was quantified and normalized to the mRNA encoding GAPDH. The relative *MX1* mRNA levels were highest in SARS-CoV-2 infected lungs without treatment. Levels of the same mRNA were reduced by treatment with Molnupiravir or STP938, as well as by the combination. However, STP938 monotherapy had the strongest impact, reducing *MX1* mRNA levels to baseline. E. Mx1 protein immunoreactivity of lung tissues, determined by immunohistochemistry (IHC). All SARS-CoV-2 infected groups, except the STP938 treated group, showed an increased Mx1 protein immunoreactivity compared to the uninfected control group. STP938 treatment alone decreased the score of Mx1 protein reactivity significantly compared to all other infected groups. Statistical evaluation was performed by the Kruskal-Wallis test with Benjamini Hochberg correction for multiple comparison (FDR 0.05). The Molnupiravir treated group and the combined treated group showed immunoreactivity comparable to the non-treated SARS-CoV-2 group. For representative images of hamster tissues, see Suppl. Fig. 6.

Taken together, these results strongly suggest that CTPSis display remarkable therapeutic efficacy against SARS-CoV-2 replication and the correlated tissue pathology. While we did not observe a synergistic effect of Molnupiravir and STP938 at the doses applied in our animal study, our results demonstrate that the CTPS inhibitor alone reduces viral load and tissue damage in an animal model of COVID-19.

## DISCUSSION

Our results indicate that CTPS inhibition fortifies the efficacy of NHC in cell culture models. Given the suboptimal performance of all currently available drugs against COVID-19, this raises the perspective of using Molnupiravir in combination with CTPSis for improved antiviral therapy. Although Molnupiravir is hardly used for COVID therapy at present, the possibility of future pandemics involving coronaviruses along with the broad activity of the drug still make its exploration worthwhile for pandemic preparedness. The same is true for the use of CTPS inhibitors as antivirals, especially in the context of its strong in vivo activity.

Our previous work indicated a similar synergy of Molnupiravir/NHC with inhibitors of DHODH (Stegmann et al., 2022), an enzyme required for the synthesis of orotate, which is then turned into orotate monophosphate and uridine monophosphate. Thus, DHODH inhibition can be compensated by the uptake and subsequent phosphorylation of uridine. Given the high uridine serum concentrations, this mechanism provides at least one explanation for the somewhat disappointing data regarding a possible synergy of these two drugs in a preclinical animal model (Stegmann et al., 2022). CTPS inhibition may at least partially overcome this problem. Even when uridine is available to the cell, RNA synthesis still requires its turnover into cytidine nucleotides, and CTPS is essential for this step. Thus, when CTPS is blocked by inhibitors in vivo, the resulting lack of cytidine nucleotides cannot be compensated by uridine uptake.

The results of our animal studies argue that CTPS inhibition strongly diminishes virus replication. There, however, Molnupiravir did not provide a detectable additional benefit. It is possible that the CTPSi concentration was too high to see any additional impact of Molnupiravir, simply because the virus was already close to full elimination with the CTPSi alone. Moreover, Molnupiravir may be metabolised to produce cytidine in the animal model, by removing the hydroxyl group at N4. If true, this would explain why Molnupiravir appeared to counteract rather than fortify CTPSis in vivo. A recent report describes the induction of interferon signaling by depleting CTPS1, since CTPS1 was found capable of deamidating and thus inactivating IRF3 (Rao et al., 2025). However, in our system, CTPS inhibition decreased the expression of the interferon-responsive *MX1* gene in the lungs of infected animals, which appears in contrast with the expected impact on IRF3 and in any case does not provide an explanation for the antiviral effects of CTPSis that we observed in vivo. Of note, however, we cannot rule out that an early interferon response driven by CTPSis might impair virus propagation, whereas the reduction in *MX1* expression at the time of autopsy might be an indirect result of virus clearance. Furthermore, it remains to be determined whether Molnupiravir or other cytidine analogues might fortify the antiviral effect of CTPSis at different dose ratios. In any case, however, our results reveal a strong antiviral activity of CTPSis in vivo.

The CTPSi CPEC was previously investigated as a cancer drug (Schimmel et al., 2007). Upon triphosphorylation, CPEC binds and inhibits CTPS activity (Kang et al., 1989). Depleting cytidine nucleotides by CPEC was found to slow down the proliferation of cancer cells and to induce cell death (Moyer et al., 1986). For this purpose, the combination of the CTPSi with cytotoxic cytosine analogues was also considered and tested using in vitro systems. Indeed, combining CPEC with gemcitabine (Verschuur et al., 2004) or cytosine-arabinoside (Ara-C) (Bierau et al., 2003) led to extensive tumor cell death. It is highly plausible that the lack of available cytidine nucleotides (through CTPS inhibition) increased the incorporation of false nucleotides into nascent DNA, explaining the cooperative drug efficacy. Thus, similar principles are applicable to antiviral and anti-cancer therapy by nucleoside analogues. Likewise, a combination of CTPSi and the cytidine analogue Lamivudine (2’-desoxy-3’-thiacytidine, 3TC) was found effective against the replication of Human Immunodeficiency Virus (Dereuddre-Bosquet et al., 2004).

In the clinics, studies involving intravenous infusion of CPEC were discontinued due to severely reduced blood pressure (hypotension) observed in some patients (Politi et al., 1995). It is unknown whether this effect was due to on- or off-target mechanisms, but the latter is made plausible by the fact that such toxicities had not been observed in animal models. Moreover, the serum concentrations of CPEC that were associated with hypotension (up to 3 µM) considerably exceeded the concentrations required for synergy with NHC in our assays (300 nM). This argues that, even in the case of on-target toxicities, there is a sufficient therapeutic window to combine CTPSis with nucleoside analogues in a clinical setting.

There are two isoforms of CTPS encoded by human and other mammalian genomes. Little biochemical differences between CTPS 1 and 2 activities are known, but their expression patterns differ considerably. CTPS2 is ubiquitously expressed, with few alterations between tissues. In contrast, CTPS1 is constitutively found in the liver and salivary glands, according to the proteinatlas.org entries (Uhlen et al., 2017). More importantly, however, CTPS1 is strongly expressed in activated T and B lymphocytes, and it is essential for their proliferation (Martin et al., 2020; Martin et al., 2014). Patients lacking properly functioning CTPS1 still develop normally, except for severe immunodeficiency. This argues in favor of using a specific CTPS1 inhibitor for immunosuppression. The question remains, however, why a specific CTPS1 inhibitor, such as STP938, was still capable of interfering with virus replication outside lymphocytes in our experiments. CTPS1 was reported to support virus replication by de-amidation of IRF3 (Rao et al., 2025), but if this would apply to the hamster model studied here, enhanced interferon signaling would be expected in response to CTPSi treatment – in contrast, we observed reduced *MX1* expression with CTPS inhibition. Perhaps more conceivably, virus infection might activate CTPS1 in the animals, e.g. through the formation of filaments termed cytoophidia (Carcamo et al., 2011; Keppeke et al., 2015; Yin et al., 2024), enhancing the contribution of CTPS1 to CTP synthesis in infected cells. This might explain the antiviral activity of CTPSis.

Rather than inhibiting CTPS, it was previously proposed to combine DHODH inhibitors with drugs that interfere with the pyrimidine salvage pathway (Liu et al., 2020), i.e. with enzymes that phosphorylate uridine or cytidine. For full antiviral activity, this would then need to be combined with an antiviral pyrimidine analogue for incorporation during RNA replication. Although requiring three inhibitory molecules together, such an approach might still represent a viable extension of the strategy outlined here.

CTPS inhibition conceivably compromises the immune response, as strongly suggested by the severe immunodeficiency of patients who carry CTPS1 mutations (Martin et al., 2020; Martin et al., 2014). At first glance, it seems counterintuitive to suppress the immune response in virus-infected patients. However, exceedingly strong inflammatory reactions (“cytokine storm”) form a considerable part of COVID-19 pathogenesis (McGonagle et al., 2021; Vora et al., 2021), and this might well be attenuated by interfering with nucleotide biosynthesis. Thus, on top of enhancing the impact of Molnupiravir, CTPS inhibition might ameliorate COVID-19 symptoms by taming excessive immune reaction and inflammation.

Nucleoside analogues, including cytidine-derived compounds, form a substantial class of antiviral agents. Molnupiravir was not only used against SARS-CoV-2, but also for treating influenza virus (Toots et al., 2019) and even Ebola virus (Bluemling et al., 2023) infections, at least in cell culture models. Other examples of antiviral cytidine analogues include 5-azacytidine, Zalcitabine, Lamivudine and Decitabine against HIV, Balapiravir against HCV, and troxacitabine against HBV (Kataev and Garifullin, 2021). Thus, CTPS inhibition has the potential of augmenting the efficacy of antiviral drug regimens in a broad range of infections.

## ACKNOWLEDGEMENTS

We thank Otto Haller and Georg Kochs, University of Freiburg, Germany, for generous gifts of antibodies to Mx1 and helpful advice on their use.

KMS and HF were members of the Göttingen Graduate School GGNB during this work.

This work was funded by the COVID-19 Forschungsnetzwerk Niedersachsen (COFONI) to MD, AV and MvK-B, by the VolkswagenStiftung AZ 9A827 to MD as well as AZ 9B785 to MD and AB-B.

## CONFLICT OF INTEREST STATEMENT

KMS, AD, HLF and MD are employees of University Medical Center Göttingen, which has filed a patent application covering the combination of inhibitors of pyrimidine synthesis and nucleoside analogues to treat viral infections (inventors: MD, KMS, AD). PEB and AEP are employees of Step-Pharma, a company addressing the use of CTPSis. The other authors declare no conflict of interest.

## SUPPLEMENTAL FIGURES

**Supplemental Figure 1:**
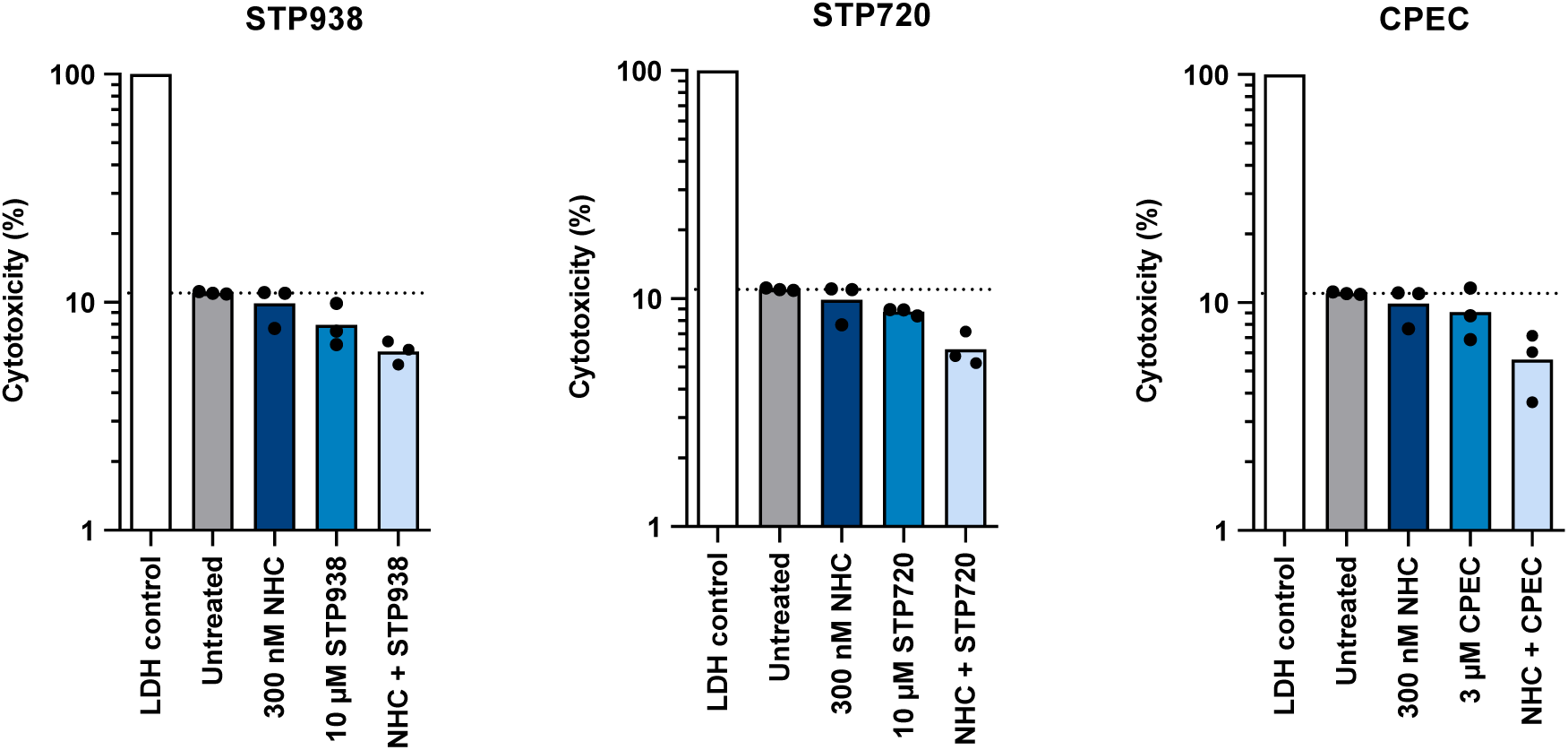
Analysis of drug cytotoxicity. Lack of overt cytotoxicity, based on LDH release, when applying NHC with higher doses of CTPSis. Upon treatment of Vero E6 cells with the drugs at the indicated concentrations, lactate dehydrogenase (LDH) in the supernatant of drug-treated cells was quantified as described in the legend to Fig. 1C, with no increase in response to drug treatment.

**Supplemental Figure 2:**
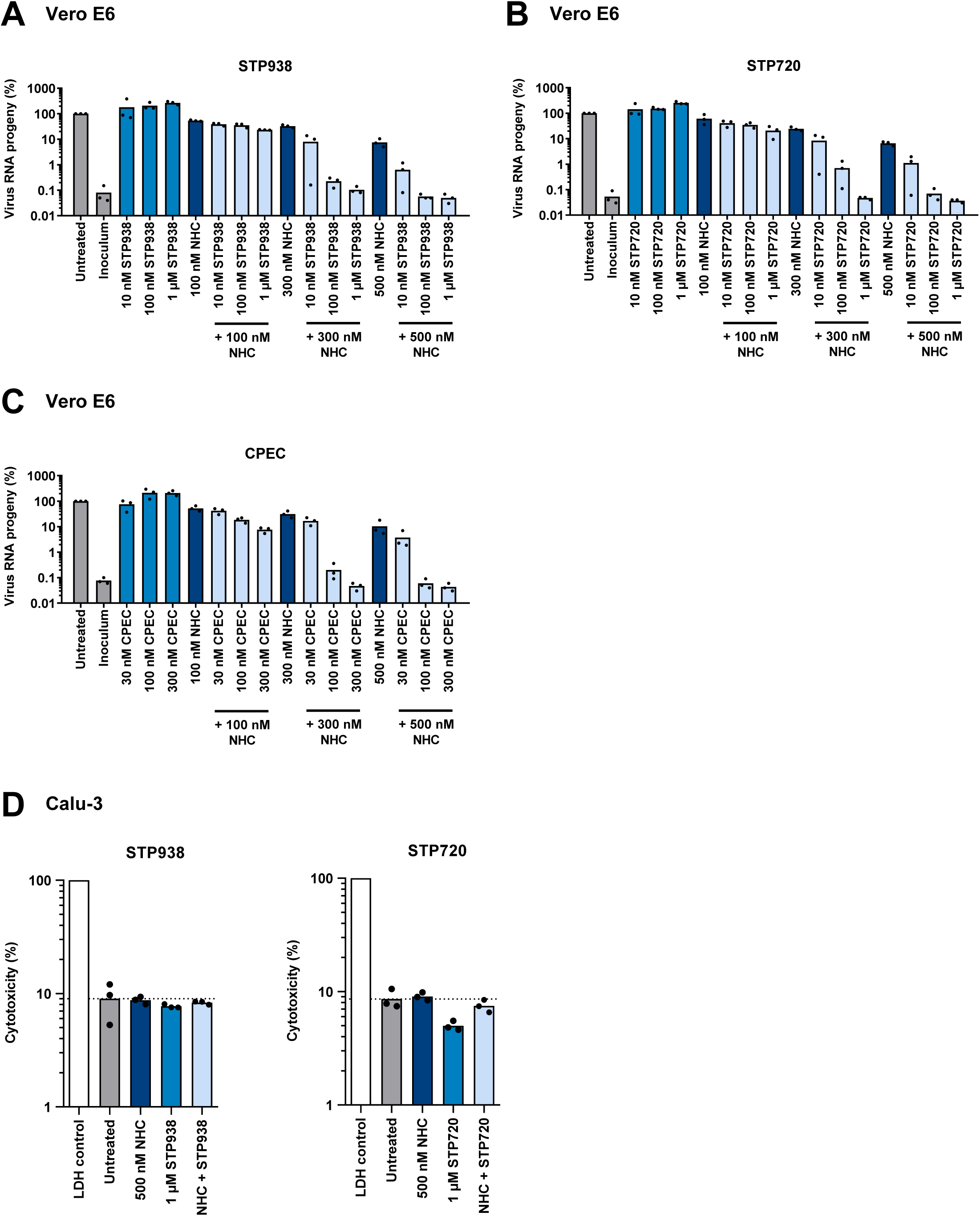
Detailed quantification of virus RNA progeny and cytotoxicity. (A-C) Data from the experiments displayed in Fig. 2A, analyzing the response of SARS-CoV-2 replication in Vero E6 cells to the indicated drugs, now displaying each measurement as a dot. For p-values, see Suppl. Table 1. (D) Lack of overt cytotoxicity when treating Calu-3 cells with drugs, as determined by the release of LDH as in Fig. 1C, corresponding to the analysis of virus RNA progeny in Fig. 2C.

**Supplemental Figure 3:**
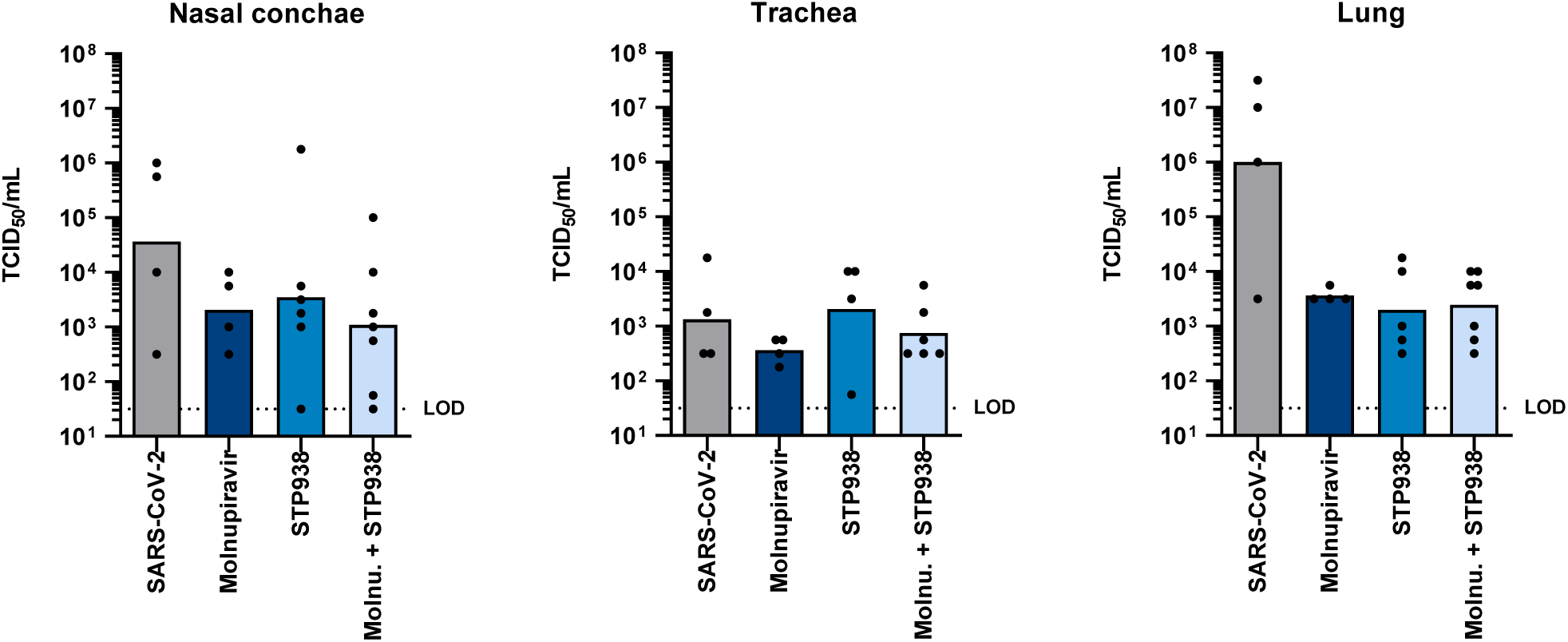
Infectious particles in hamster tissues at post mortem analysis. The amount of infectious particles was determined by TCID_50_ measurements at day 6 after infection. At this time, only lung tissue displayed higher amounts of virus in infected but untreated hamsters *vs*. drug-treated animals.

**Supplemental Figure 4:**
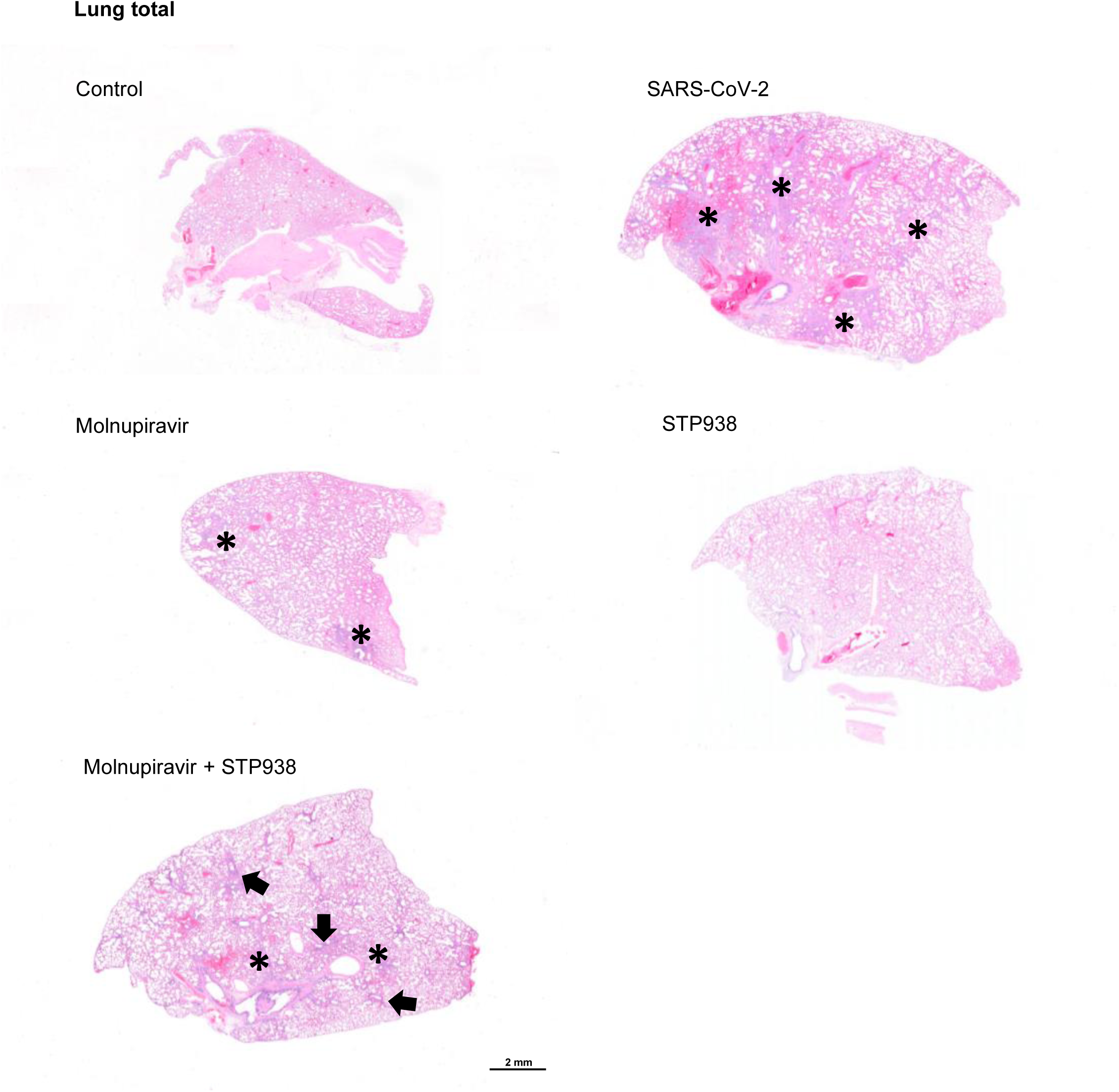
Representative images of hamster lung tissue (overview) Upon treatment and infection of hamsters (Fig. 6A), the lungs of the animals were subjected to analysis of pathological alterations upon H&E staining. The results summarized in Fig. 6C are shown by depicting representative sections of left lung lobes. While lungs of the non-infected control hamsters showed minimal edema and/or bleeding, multifocal consolidation, alveolitis and atelectasis (asterisks) was present in the SARS-CoV-2 infected hamsters without treatment. Lesions were nearly completely resolved in the STP938 treated group, while both groups with Molnupiravir showed some improvement of lung pathology but no resolution. In the Molnupiravir and STP938 group, lesions were accentuated in the perivascular areas (arrows).

**Supplemental Figure 5:**
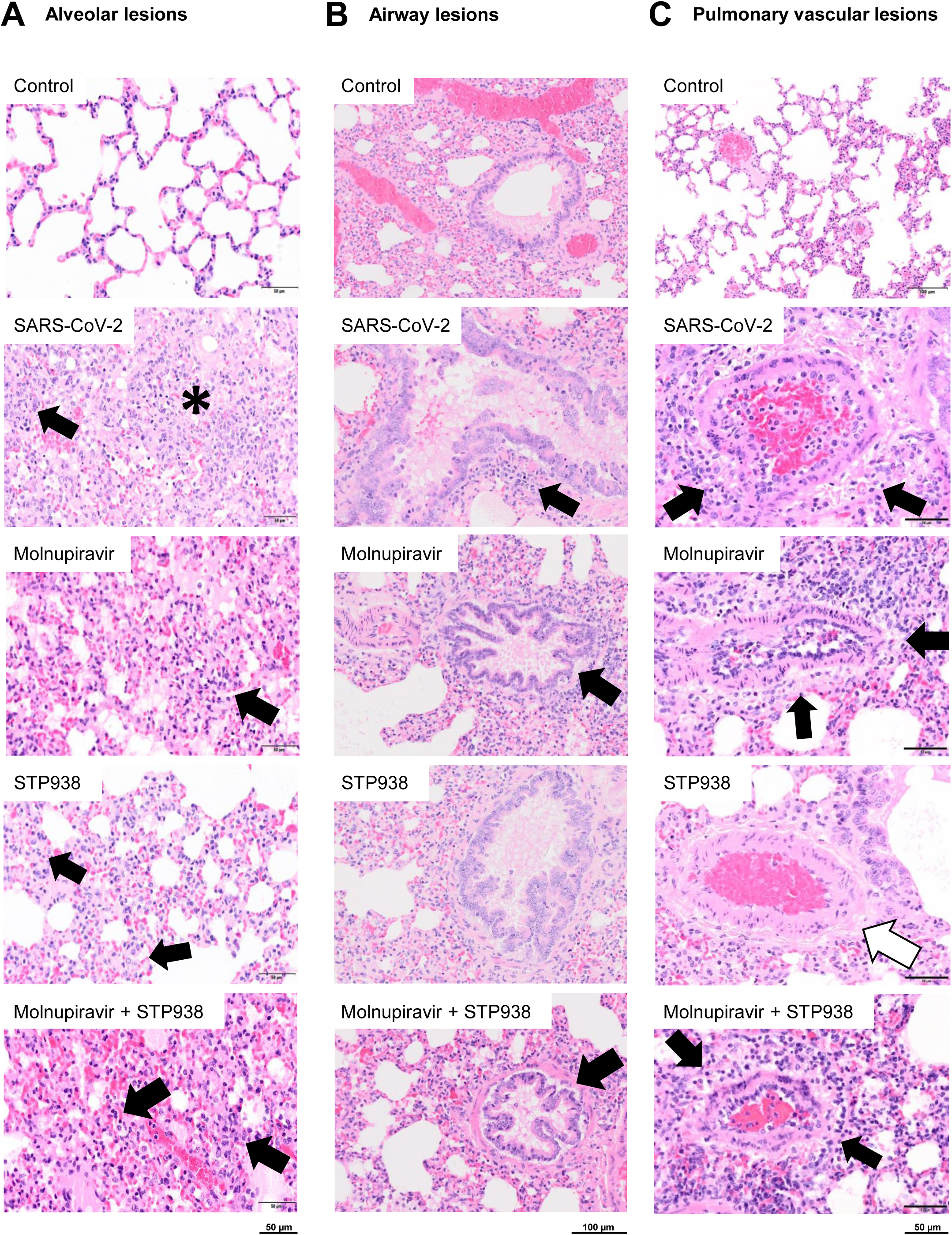
Images of hamster lung tissue (details) The results summarized in Fig. 6C are depicted, showing representative details of the lungs as in Suppl. Fig. 4. (A) Alveolar lesions: While inflammatory lesions in the non-infected control group were absent, marked multifocal consolidation of lung parenchyma with proliferation of type II pneumocytes (asterisk) was notable in the untreated SARS-CoV-2 infected hamsters. Consolidation was less pronounced in all treatment groups with STP938 only treatment revealing mostly edema and mild alveolitis. Alveolitis was characterized by intraalveolar infiltration of heterophils and macrophages (black arrows), occasional pneumocyte necrosis, rare presence of multinucleated pneumocytes and alveolar edema and fibrin deposition. (B) Airway lesions: While inflammatory lesions in the non-infected control group were absent, mild to moderate multifocal bronchitis with peribronchial infiltration (arrows) was present in the untreated SARS-CoV-2 infected hamsters. Inflammation was less pronounced in all treatment groups with STP938 only treatment, revealing only minimal to absent lesions. Bar: 100µm (C) Pulmonary vascular lesions: In SARS-CoV-2 infected untreated and Molnupiravir-treated hamsters, pronounced vascular lesions were noted. Lesions consisted of perivascular edema and mononuclear infiltration (black arrows). In addition, marked hypertrophy of endothelial cells was noted. In the STP938-treated hamsters, vascular lesions were nearly resolved with mild perivascular edema as the only detectable lesion (white arrow). The combined treatment led to decreased severity of vascular lesions. No lesions were detectable in the non-infected and untreated control group.

**Supplemental Figure 6:**
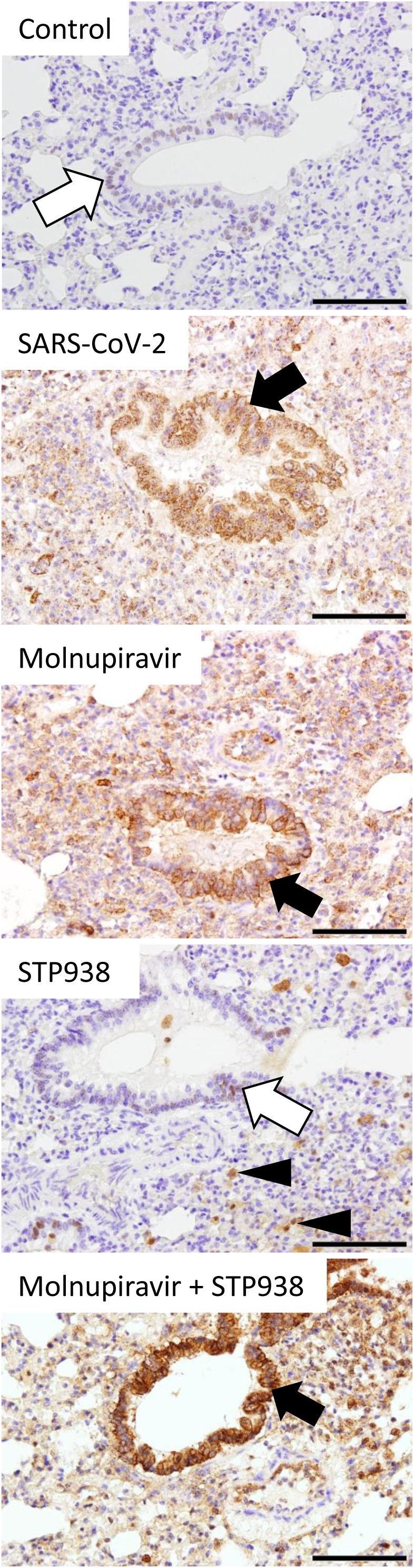
Immunohistochemical analysis of Mx protein expression in hamster lung tissue. Comparison of Mx protein immunolabeling, corresponding to the quantification in Fig. 6D: In non-infected hamsters, only minimal immunolabeling was present, while it was markedly increased in SARS-CoV-2 infected, untreated hamsters. A similar immunoreactivity was present in the Molnupiravir treated hamster and in the combination treatment group. Immunolabeling in these three groups was present in the cytoplasm of bronchiolar epithelium, vascular endothelial cells and scattered immune cells/type II pneumocytes (black arrows). Only mild labeling was observed in the STP938 treated hamsters. In the uninfected and in the STP938 treatment group, scattered bronchiolar immunolabeling was nuclear (white arrows). In the STP938 group, scattered leukocytes and type II pneumocytes showed nuclear to cytoplasmic immunolabeling (arrowheads). Bar: 100µm

**Supplemental Table 1:** p-values of the data displayed in Fig. 2A and Suppl. Fig. 2A-C. Statistical analysis was conducted using GraphPad Prism 9. A two-sided unpaired Student’s t-test was applied, with significance defined at p ≤ 0.05.

## MATERIALS AND METHODS

## RESOURCE AVAILABILITY

### Lead contact

Further information and requests for resources and reagents should be directed to and will be fulfilled by the lead contact, Matthias Dobbelstein.

### Materials availability

This study did not generate new unique reagents.

### Data and code availability

- All data reported in this paper will be shared by the lead contact upon request.
- This paper does not report original code.
- Any additional information required to reanalyze the data reported in this paper is available from the lead contact upon request.

## EXPERIMENTAL MODEL AND SUBJECT DETAILS, METHOD DETAILS AND QUANTIFICATION AND STATISTICAL ANALYSIS

These sections are combined for better readability, since the methods and analyses were different for each experimental model system.

### SARS-CoV-2 replication in Vero E6 and Calu-3 cells

#### Cell culture

Vero E6 cells (Vero C1008) were cultured in Dulbecco’s modified Eagle’s medium (DMEM with GlutaMAX^TM^, Gibco), supplemented with 10% fetal bovine serum (Merck), 50 units/mL penicillin, 50 μg/mL streptomycin (Gibco), 2 µg/mL tetracycline (Sigma) and 10 µg/mL ciprofloxacin (Bayer). Cultures were maintained at 37°C in a humidified environment with 5% CO_2_. Calu-3 cells were cultured in Eagle’s Minimum Essential Medium (EMEM, ATCC) supplemented with 10% fetal bovine serum and penicillin/streptomycin.

#### Treatments and SARS-CoV-2 infection

20,000 cells per well were plated in 24-well-plates using medium containing 2% fetal bovine serum (FBS) and incubated for 8 hrs at 37 °C. The cells were then treated with β-D-N^4^-Hydroxycytidine (NHC/EIDD-1931, Cayman Chemical 9002958) and/or the CTPSis cyclopentenyl cytosine (CPEC, Biosynth NC58281), STP938 (kindly provided by StepPharma), STP720 (kindly provided by StepPharma) as well as to uridine (Selleckchem S2029) or cytidine (Selleckchem S2053) at the concentrations indicated in the figures. Following this treatment, cells were infected with SARS-CoV-2 at a multiplicity of infection (MOI) of 0.1 and incubated for 48 hrs at 37 °C, as previously described (Stegmann et al., 2021). The original SARS-CoV-2 ‘wildtype’ was isolated from a patient sample obtained in March 2020 in Göttingen, Germany, with the exact genome sequence published previously (Stegmann et al., 2021). The SARS CoV-2 Omicron variant (BA.5, EVA: 009V-05261) was kindly provided by the Robert Koch Institute, Berlin, Germany.

#### TCID_50_ determination

To calculate the Median Tissue Culture Infectious Dose (TCID_50_) per mL in cell culture supernatants, 20,000 Vero E6 cells per well were treated and infected as described above. At 48 hrs post infection (p.i.), the virus-containing supernatant was titrated to calculate the TCID_50_/mL. To this end, Vero E6 cells were incubated with ten-fold dilutions (quadruplicates) of virus for another 48 hrs. Cells were then fixed with 4% formaldehyde in PBS for 1 hour at room temperature. After permeabilization with 0.5% Triton X-100 in PBS for 30 min and blocking in 10% FBS/PBS for 10 min, the SARS-CoV-2 nucleoprotein (N) was visualized via immunofluorescence staining (primary antibody: Sino Biological #40143-R019, 1:8000; secondary antibody: Alexa Fluor 546 donkey anti-rabbit IgG, Invitrogen, 1:500). Fluorescence signals were detected by automated microscopy using a Celígo^®^ Imaging cytometer.

For the calculation of the TCID_50_ titer in swab or nasal lavage samples as well as organ homogenates, samples were serially diluted in MEM containing 2% FCS and 100 Units Penicillin / 0.1 mg Streptomycin (P/S) (Millipore Sigma, Germany). Vero E6 cells were incubated with 100 µl of ten-fold dilutions of sample dilutions added in quadruplicates for 1 hr at 37°C before 100 µl MEM containing 2% FCS and P/S were added per well and plates were incubated for 5 days at 37°C and 5% CO_2_. Supernatant was removed and cells were fixed with 4% formalin. Next, plates were stained with 1% crystal violet and wells showing a cytopathogenic effect were counted.

In both cases, the viral titer was determined according to Spearman and Kärber (Kärber, 1931).

#### Quantitative RT-PCR for virus RNA quantification

To extract RNA, the SARS-CoV-2-containing cell culture supernatant was mixed in a 1:1 ratio with the Lysis Binding Buffer from the MagNA Pure LC Total Nucleic Acid Isolation Kit (Roche) to inactivate the virus. Viral RNA was isolated using Trizol LS, chloroform, and isopropanol. Following an ethanol wash of the RNA pellet, the isolated RNA was reconstituted in nuclease-free water.

Total RNA from swab, nasal lavage samples and tissue homogenates was extracted as described earlier (Schlottau et al., 2020). SARS-CoV-2 RNA was detected (Cormann et al., 2020). Quantitative Realtime PCR was performed with the qScript XLT One-Step RT-qPCR ToughMix (QuantaBio/VWR). Viral genome copy numbers were calculated from standard curves determined for 10^-2^ to 10^-5^ dilutions containing known copy numbers of SARS-CoV-2 (Blaurock et al., 2022).

Quantitative RT-PCR, based on a previously established assay utilizing a TaqMan probe (Corman et al., 2020), was conducted to measure virus RNA yield. Specific oligonucleotides amplifying a genomic region corresponding to the envelope protein gene (26,141 – 26,253), were used for qRT-PCR.

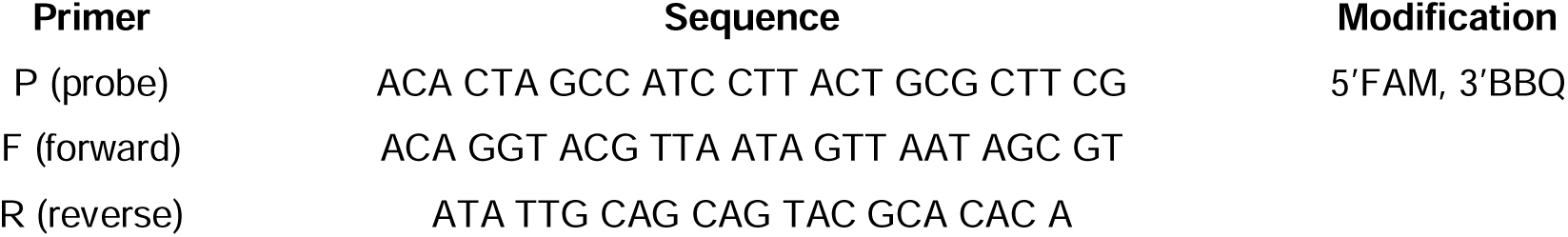

The data acquired via qRT-PCR were presented as virus RNA copies per mL or virus RNA progeny (%), where the amount of SARS-CoV-2 RNA detected upon infection without any treatment was set as 100%, and the remaining RNA quantities were normalized accordingly.

#### Determination of drug synergy

Vero E6 cells were subjected to treatment and/or infection as specified, and RNA from the cell culture supernatant was isolated and quantified by qRT-PCR. The quantity of SARS-CoV-2 RNA detected upon infection without any treatment was set to 100% virus yield (0% inhibition). The remaining samples were quantified as a percentage of the control. The Bliss independence model (Bliss, 1939), computed using the synergyfinder website (Ianevski et al., 2020), was utilized to assess synergy between drug combinations.

#### Immunofluorescence analyses

5,000 Vero E6 cells per well were seeded onto 8-well chamber slides (Nunc) and were treated/infected as indicated. Following 48 hrs of SARS-CoV-2 infection, cells underwent a PBS wash and were fixed with 4% formaldehyde in PBS for 1 hr at room temperature. Subsequently, cells were permeabilized with 0.5% Triton X-100 in PBS for 30 min, followed by a 10-minute blocking step in 10% FBS/PBS. Primary antibodies targeting the SARS-CoV-2 Nucleoprotein (N; Sino Biological #40143-R019, 1:8000) and Spike protein (S; GeneTex#GTX 632604, 1:2000) were applied overnight. The following day, secondary antibodies, Alexa Fluor 546 donkey anti-rabbit IgG and Alexa Fluor 488 donkey anti-mouse IgG (Invitrogen, 1:500, diluted in blocking solution), were added along with 4′,6-diamidino-2-phenylindole (DAPI) for 1.5 hrs at room temperature. Slides were mounted using Fluorescence Mounting Medium (DAKO) and fluorescence signals were visualized via microscopy (Zeiss Axio Scope.A1).

#### Immunoblot analyses

Upon treatment and SARS-CoV-2 infection as described above, cells were washed twice in PBS and harvested in radioimmunoprecipitation assay (RIPA) lysis buffer, a solution consisting of 20 mM TRIS-HCl pH 7.5, 150 mM NaCl, 10 mM EDTA, 1% Triton-X 100, 1% deoxycholate salt, 0.1% SDS, 2 M urea and protease inhibitors. Samples were briefly sonicated and protein content was determined using the Pierce BCA Protein assay kit (Thermo Fisher Scientific). Once the protein quantities were adjusted, the samples were heated up to 95 °C in Laemmli buffer for 5 min and then separated through sodium dodecyl sulfate polyacrylamide gel electrophoresis (SDS-PAGE). To determine the presence of SARS-CoV-2 proteins, the separated proteins were transferred to a nitrocellulose membrane, blocked in 5% (w/v) non-fat milk in TBS containing 0.1% Tween-20 for 1 hr, and incubated with primary antibodies at 4 °C overnight, followed by a subsequent incubation with peroxidase-conjugated secondary antibodies (donkey anti-rabbit or donkey anti-mouse IgG, Jackson Immunoresearch). The presence of specific proteins including SARS-CoV-2 Spike- and Nucleoprotein, CTPS1, CTPS2 and HSC70 (loading control) was detected using either Super Signal West Femto Maximum Sensitivity Substrate (Thermo Fisher) or Immobilon Western Substrate (Millipore).

| **Antibody** | **Source (Catalogue number)** | **Dilution** |
| --- | --- | --- |
| SARS-CoV-2 Spike | GeneTex GTX 632604 | 1:1,000 |
| SARS-CoV-2 Nucleoprotein | Sino Biological 40143-R019 | 1:5,000 |
| CTPS1 | Sigma-Aldrich HPA-051322 | 1:500 |
| CTPS2 | Sigma-Aldrich HPA-017437 | 1:1,000 |
| HSC70 | Santa Cruz sc-7298 | 1:10,000 |

#### Quantification of LDH release to determine cytotoxicity

3,500 Vero E6 cells were seeded into 96-well-plates and treated with CTPSis and/or NHC as specified, for a duration of 72 hrs. The quantification of lactate dehydrogenase (LDH) release into the cell culture medium was conducted by bioluminescence using the LDH-Glo^TM^ Cytotoxicity Assay kit from Promega. To establish the maximum LDH release, untreated cells were exposed to 10% Triton X-100 for 15 min, whereas the medium background served as a negative control. Percent cytotoxicity, indicating the proportion of LDH released into the media in comparison to the total LDH content within the cells, was computed utilizing the following formula.

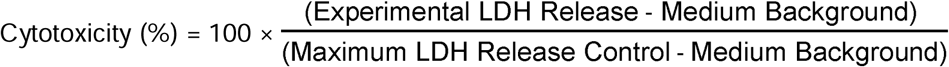

#### Statistical analysis of cell-based experiments

Statistical analyses were conducted using GraphPad Prism 9. Unless stated otherwise, two-sided unpaired Student’s t-tests were applied, with significance defined at p ≤ 0.05. Statistical significance is indicated by asterisks as follows: **** for p ≤ 0.0001; *** for p ≤ 0.005; ** for p ≤ 0.01; and * for p ≤ 0.05.

#### SARS-COV-2 treatment study in Golden Syrian hamsters

Male Golden Syrian hamsters (*Mesocricetus auratus*), 5-7 weeks old with a body weight of 80 – 100 g, were obtained from Janvier Labs, France. Three hamsters were housed in individually ventilated cages (IVC). Animals had ad libitum access to food and water. The animals’ well-being and body weight were checked daily. Handling and sampling were performed starting with the uninfected group and continuing from the low dose groups to the high dose groups to minimize the contamination risk.

#### Drug treatment

Animals received either 250 mg/kg Molnupiravir dissolved in 2% Tween 80 2%, 10% PEG400, 66% Water 66% and 22% strawberry sirup orally two times daily in a volume of 500 µl, or 50 mg/kg STP938 once daily by subcutaneous (s.c.) injection, or a combination of both. The mock infected and the positive control animals received the same volume of Tween 20 / PEG 400, Water and strawberry Sirup orally and/or PBS s.c.

#### Inoculation routes

Hamsters were inoculated orotracheally route with 100 µl containing 1*10^3^ TCID_50_. Oral swab samples were collected in DMEM containing P/S on days 1, 3 and 5 post infection, while Nasal lavage samples were collected under isoflurane anaesthesia at days 2, 4, and 6, post infection by flushing 200 µl PBS along the animal’s nose. At 6 dpi, hamsters were euthanized by deep isoflurane anaesthesia, cardiac exsanguination and cervical dislocation, and nasal conchae, trachea and lung samples were collected and stored at -80 °C for virological analysis. Tissue samples were also stored in 4% neutral-buffered formalin for histopathological analysis. Serum was separated from the collected blood.

#### Pathology, staining procedures, image acquisition and score determination

Formalin-fixed paraffin embedded left lung lobes and tracheal sections were submitted to the Department of Pathology, University of Veterinary Medicine Hannover, Hannover, Germany. Tissue blocks were trimmed and stained with hematoxylin-eosin according to routine protocols. Respiratory lesions were scored after histopathological examination by two veterinary pathologists (TS, WB) semiquantitatively as described (Aksu et al., 2024; Armando et al., 2022) with minor modifications. In summary, alveolar, conductive airways and vascular lesions were examined using a 4-tier-scoring system. Extent and severity of tracheal, airway and alveolar lesions were multiplied before summing of all scores to get to the final scores.

For statistical analysis and data visualization, GraphPad Prism software version 10 for Windows™ (GraphPad software, San Diego, California, USA) was used. Data sets were compared by Kruskal-Wallis multi-comparison test with subsequent pairwise Mann-Whitney-U-tests and Benjamini-Hochberg correction for multiple comparisons (False Detection Rate: 0.05). Exact p values ≤ 0.05 were assumed statistically significant.

SARS-CoV-2 nucleoprotein immunohistochemistry was performed according to established protocols (Armando et al., 2022). After deparaffinization, rehydration and incubation, sections were incubated with a mouse monoclonal antibody against the SARS-CoV-2 nucleoprotein (Sino Biological, Peking, China-40143-MM05). Immunolabeling was visualized using the EnVision+ polymer system (Dako Agilent Pathology Solutions) and 3,3’-diaminobenzidine tetrahydrocloride (DAB) as chromogen (Sigma Aldrich). Immunohistochemistry to detect Mx1 protein was done as previously published (Klotz and Gerhauser, 2019), with minor modifications. Briefly, sections were deparaffinized and endogenous oxidase was blocked. Afterwards, the slides were pretreated in citrate buffer using a microwave oven. Subsequently, the slides were incubated with anti-Mx1 antibody (M143, kindly provided by O. Haller and G. Kochs, University Medical Center Freiburg, Freiburg, Germany, dilution: 1:1000) overnight at 8°C. As secondary antibody, goat-anti-mouse IgG (BA-9200, Vector Laboratories Inc., CA, USA) was applied. For visualization of the immunolabeling, the Avidin-Biotin-Complex method (Vectastain Elite ABC Kit, Vector Laboratories Inc.) and DAB as chromogen were used. For negative controls, the primary antibodies were replaced by the respective protein concentration of ascitic fluid from non-immunized BALB/cJ mice.

All slides were evaluated with a Zeiss Axioscope (Zeiss, Göttingen, Germany; field of view at 400x magnification: 0.16 mm²). Whole slides were scanned with the Olympus VS200 slide scanner (Olympus Deutschland GmbH, Hamburg, Germany) and images of animals representing group median score were exported with the respective OlyVIA software.

#### Analysis of the interferon response in hamster RNA samples

##### cDNA Synthesis

RNA samples derived from hamster lungs were obtained from the Friedrich-Loeffler-Institut, Bundesforschungsinstitut für Tiergesundheit. RNA concentrations were determined using a NanoDrop Spectrophotometer DS-11+, Denovix, and the samples were adjusted to the lowest concentration. For primer annealing, a mixture of the RNA as well as dNTPs (R0182, Thermo Scientific), oligo(dT) (TTT TTT TTT TTT TTT TTT TTT TTV, Metabion) and random nanomers (NNN NNN NNN, Metabion) was brought to 70 °C for 5 min; subsequently, cDNA synthesis was performed using M-MuLV reverse transcriptase with reaction buffer (M0253S, BioLabs) and an RNase inhibitor (M0307 L, NEB) for 1 hr at 42 °C, followed by 5 min at 95 °C for termination.

##### qPCR

cDNA was quantified by qPCR using primers corresponding to hamster Mx1 (maMx1_for_2 CCGTGAATTCCCAGAACTAAGA; maMx1_rev_2 GAGGTTGTTCACAGAGTCCATAAA) and a SYBR-green master mix containing dNTPs (R0182, Thermo Scientific), Taq polymerase (1800.4, Primtech) and SYBR Green (S7567, Life Technologies). The data were normalized to hamster GAPDH (maGAPDH_for GCAGTTCAAAGGCACAGTCA; maGAPDH_rev TGGTGGTGAAGATGCCAGTA) and plotted using GraphPad Prism 10.

## Notes

### Competing Interest Statement

KMS, AD, HLF and MD are employees of University Medical Center Goettingen, which has filed a patent application covering the combination of inhibitors of pyrimidine synthesis and nucleoside analogues to treat viral infections (inventors: MD, KMS, AD). PEB and AEP are employees of Step-Pharma, a company addressing the use of CTPSis. The other authors declare no conflict of interest.

